# CG2AT2: An Enhanced Fragment-based approach for Serial Multi-scale Molecular Dynamics simulations

**DOI:** 10.1101/2021.03.25.437005

**Authors:** Owen N. Vickery, Phillip J. Stansfeld

## Abstract

Coarse-grained molecular dynamics provides a means for simulating the assembly and interactions of macromolecular complexes at a reduced level of representation, thereby allowing both longer timescale and larger sized simulations. Here, we describe an enhanced fragment-based protocol for converting macromolecular complexes from coarse-grained to atomistic resolution, for further refinement and analysis. While the focus is upon systems that comprise an integral membrane protein embedded in a phospholipid bilayer, the technique is also suitable for e.g. membrane-anchored and soluble protein/nucleotide complexes. Overall, this provides a method for generating an accurate and well equilibrated atomic-level description of a macromolecular complex. The approach is evaluated using a diverse test set of eleven system configurations of vary size and complexity. Simulations are assessed in terms of protein stereochemistry, conformational drift, lipid/protein interactions, and lipid dynamics.

## Introduction

Membrane proteins are fundamental to life. They are the gate-keepers into and out of a cell and critical receptors for transmitting signals across cellular membranes. In the majority of organisms they comprise ~25% of genes, and from a pharmaceutical perspective they form the targets for roughly half of all drugs^1–3^. Structural determination of these proteins has advanced considerably in the past decade, with improvements to the protocols for X-ray diffraction and the advent of atomic-resolution cryo-Electron Microscopy (cryo-EM) resulting in a total over 5,000 membrane protein structures, of which there are more than 1,000 unique proteins (see http://memprotmd.bioch.ox.ac.uk/stats for the most recent data)^4^. Despite these improvements the lipids in the surrounding environment are frequently difficult to capture, either being lost in the purification process, too diffuse to accurately determine atomic coordinates^5^, or difficult to assign as a defined lipid type.

Molecular dynamics (MD) simulations provide an ideal tool for capturing the lipid environment around these membrane protein structures, as we have shown through our database of all membrane embedded structures, MemProtMD^4^. The use of coarse-grained (CG) modelling allows for the observation of slow processes such as membrane bending and deformations^6^, protein-lipid interactions^7,8^ and association of soluble proteins to bilayers^9^. The simulation time required for these events is generally beyond that afforded to conventional atomistic simulations^10^. CG-MD simulations enable a well-configured springboard for studying, e.g., molecular and ionic transport, ligand binding, structural dynamics, and lipid interactions of proteins^10,11^. However, first one must translate the system from a CG representation to an atomic-level description. A number of methods have been developed for the conversion of a CG system to atomistic, including, e.g., a simulated annealing methodology^12^, a geometric-based *Backward* protocol^13^, machine learning algorithms^14,15^ and a previous iteration of the fragment-based *CG2AT* approach^16^. In all cases, the methodologies permit serial multi-scale MD simulations^17^.

Here we describe a complete reworking and enhancement of our fragment-based approach, introducing an assortment of new features, an increased level of flexibility, enhanced scalability and greatly improved interoperability. We apply our method to a range of molecular systems, from biological membranes, to integral membrane proteins, lipid-modified lipoproteins and DNA-bound soluble proteins.

## Methods

### Implementation and Database

To enable the conversion of the CG system to an atomistic representation, the stereochemistry of each bead and molecule must be reintroduced. Various conversion methods have been previously proposed to enable conversion from a CG system in a *de novo* manner (see above). For CG2AT2, we implement a similar fragment-based methodology to the previous iteration of CG2AT, in which small atomistic fragments containing all the chemical and stereochemical information required are rigidly aligned to each CG bead.

For CG2AT2 we have assembled a database of fragment files that are related to the commonly used force fields, e.g. CHARMM^18^, AMBER^19^, GROMOS^20^ and OPLS^21^. To simplify the development and usability of the fragment database, minimal information is required beyond the atomistic fragments (Fig. 1A) and an optional itp file (e.g. POPC.itp).

**Figure 1:**
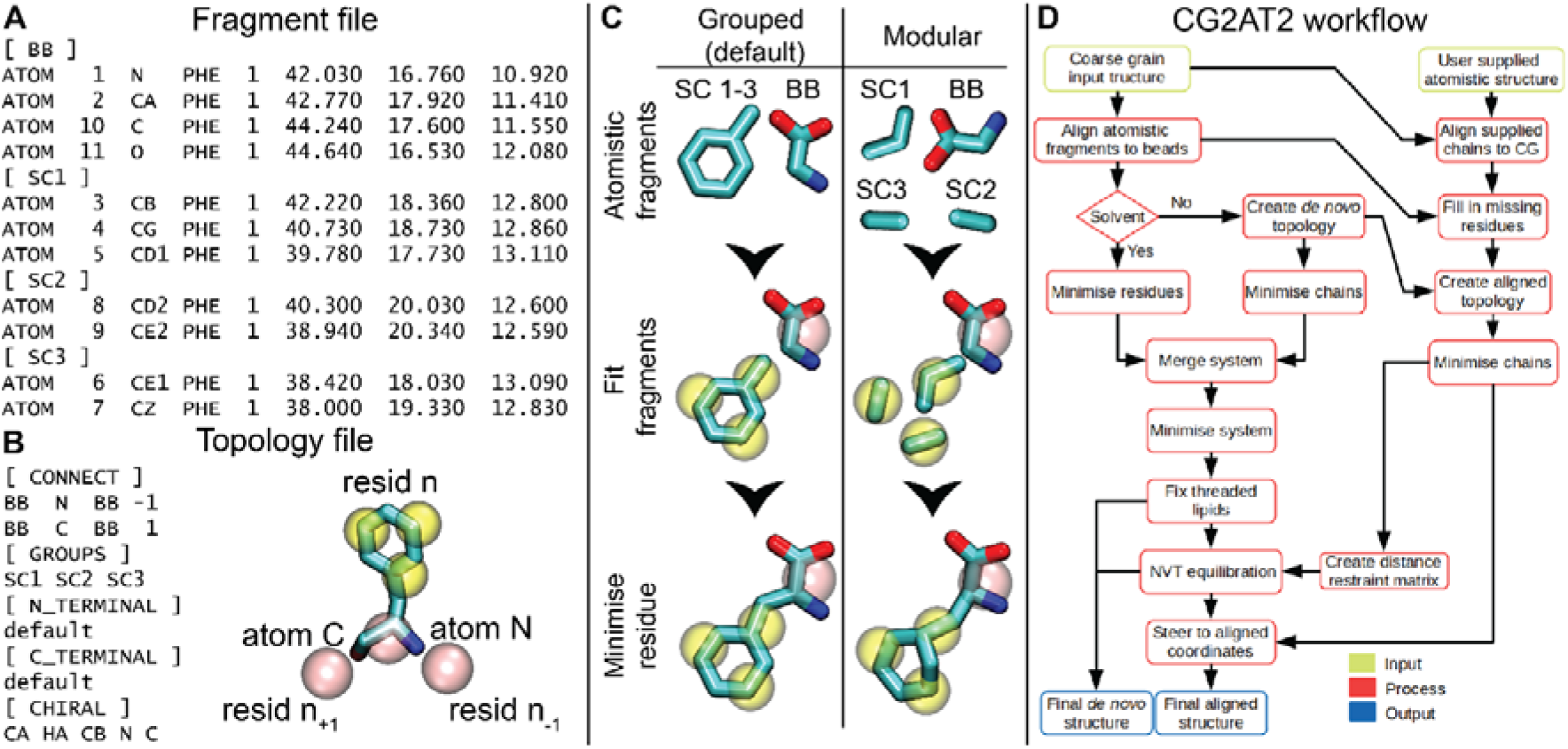
Workflow and database composition. (**A**) The fragment file for phenylalanine. Each fragment has a header containing the corresponding CG bead name, e.g. [BB], followed by the atoms within the fragment. If the molecule is not processed by `pdb2gmx` then the atom numbers should match the supplied topology itp file. (**B**) The topology file for phenylalanine. There are six sections within the topology file: `CONNECT` details the connectivity of the backbone and should follow a four column convention: bead name, associated atom name, connected bead name and numeric direction of connected bead. `GROUPS` contain the fragment groupings as shown in panel C. `N_TERMINAL` and `C_TERMINAL` contain any non-standard atomistic termini for the residue. `HIRAL` contains any stereochemistry information present within the residue. This is described in the column format: central atom, atom to move, atom 1, atom 2 and atom 3. (**C**) Graphical depiction of the grouped (left) and modular (right) grouping methodologies implemented within CG2AT2. The conversion of phenylalanine is shown. Here the grouped approach improves the conversion quality. (**D**) Description of the conversion of CG to AT. Initially CG2AT2 converts the CG system in a *de novo* manner from the fragments supplied. If an atomistic protein structure is provided, this structure is aligned with the CG coordinates, with the *de novo* protein coordinates steered to the aligned protein coordinates to remove clashes with surrounding atoms, e.g. lipids.

The fragment files are stored within a single coordinate file, where the residue is separated by bead names into fragments (Fig. 1A). The variable lengths and protonation states of multimeric macromolecules such as proteins, necessitates that the topologies are generated on the fly by the GROMACS^22^ tool `pdb2gmx`. However, individual non-protein molecules. e.g. palmitoyl-oleoyl-phosphatidylcholine (POPC), can use the supplied topology itp file in the database.

Additional information which cannot be retrieved from the itp and/or force field files is supplied in a topology file. This topology file contains four sections to aid the conversion. This file details: inter-residue connectivity, fragment groupings, terminal residue information and chiral groups (Fig. 1B). The chiral groups section enables the appropriate stereochemistry to be adhered to upon conversion. This can be aided further by grouping atomistic fragments together, so that they are structurally aligned as a single unit to the CG beads (Fig. 1C). For polymers, i.e. proteins, sugars and nucleotides, inter-residue connectivity makes sure residues are appropriately linked, and the terminal residue states options applies appropriate charge states to the first and last residues.

An optional position restraint file for the non-protein residues can also be included. These restraints may be applied during the equilibration and steering steps to minimise deviation from the CG system.

### Conversion protocol

The conversion protocol of CG2AT2 can be condensed into four core steps: conversion, minimisation, integration, and equilibration (Fig. 1D).

#### Conversion

The initial conversion phase consists of atomistic fragment fitting to the CG beads. Here, the atomistic fragments are aligned by their centre-of-mass (COM) to their respective CG bead. The aligned coordinates are then rotated to minimise bond distances with their adjacent connected beads. In this manner, each fragment can be treated individually with no knowledge of the surrounding atomistic fragments (Fig. 2A). To improve upon the initial conversion, we have implemented a fragment grouping system, detailed in an optional topology file (Fig. 1B), which allows for multiple fragments to be treated as a single fragment e.g., an amino acid side chain (Fig. 1C).

**Figure 2:**
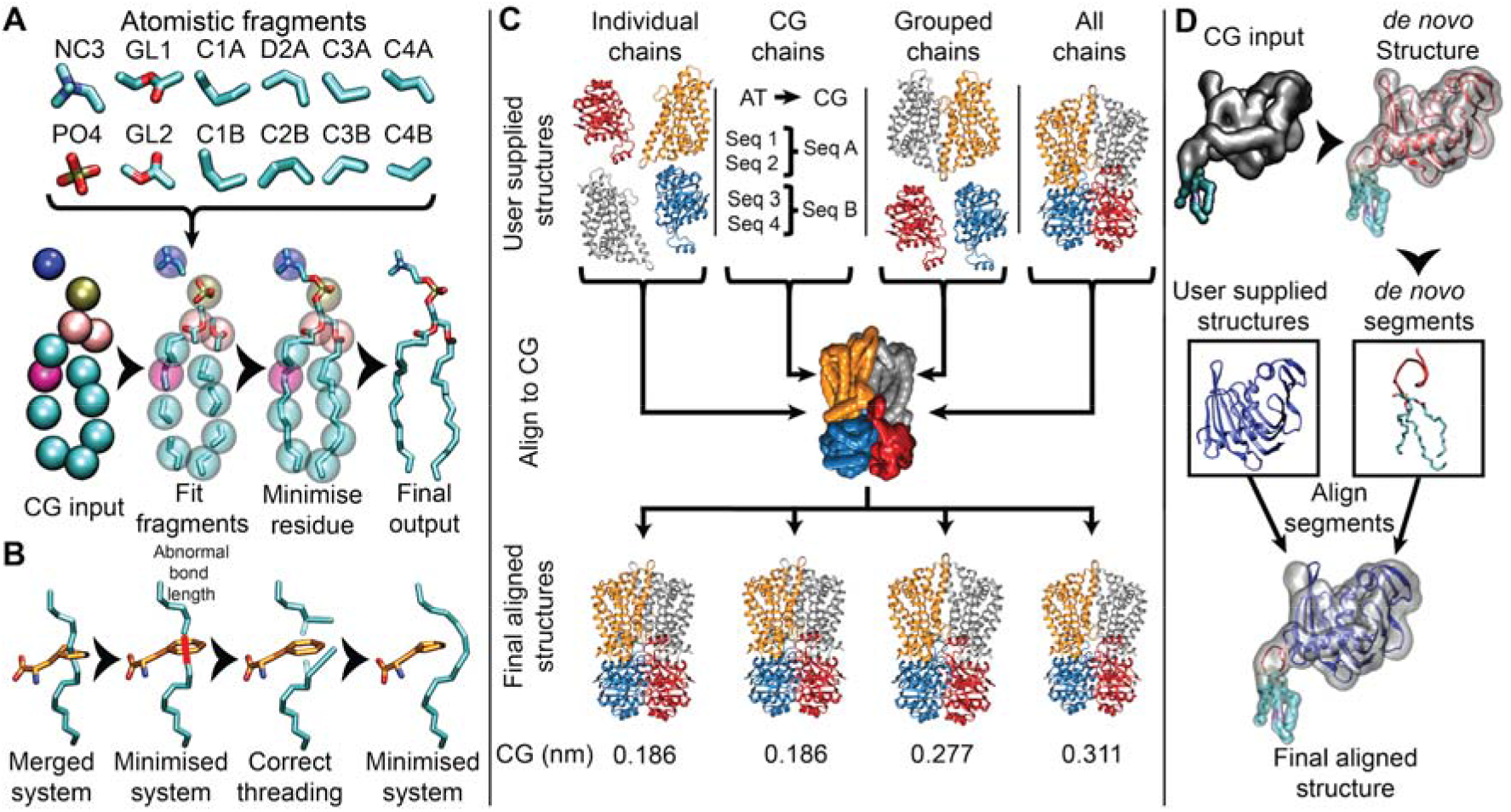
A graphical depiction of the conversion of Martini’s POPC to CHARMM36 atomistic representation. (**A**) The general conversion protocol implemented with CG2AT2. Here the individual fragments are aligned within each bead and rotated to minimise the distance between connecting beads. Once aligned, the residue undergoes the first round of minimisation. Hydrogens are hidden for clarity. (**B**) The disentanglement of a lipid tail from the sidechain of phenylalanine. If an abnormally long bond is detected after the initial minimisation of the merged system, the two non-hydrogen atoms and connected hydrogens are translated on a vector away from the threaded amino acid backbone. The system is then minimised to correct the bond lengths between the adjusted atoms. (**C**) A depiction of alignments available for the conversion of CG to AT. Here, the user is able to select whether to align the atomistic chains individually, grouped by CG chain, in selective groups or as a single macromolecular unit. RMSD measurements are highlighted: Cα against CG BB. (**D**) Flow chart of the alignment of individually supplied atomistic chains. Here, LolB is created in the *de novo* manner depicted in the red cartoon. The supplied atomistic chains (coloured blue) are aligned to the CG BB beads. The missing residues highlighted during chain alignment are copied from the *de novo* structure and mapped onto the aligned structure.

#### Minimisation

After a successful conversion, each molecule type undergoes a single round of minimisation by the steepest descent algorithm. This corrects the majority of bonds lengths, angles and any steric clashes.

#### Integration

Following a successful build of each molecule type, the system components are reintegrated and minimised. In rare cases upon reintegration, it is possible for lipid tails to be threaded through aromatic structures such as phenylalanine. To detect these instances, abnormal bond lengths are used as an indicator of a threaded molecule. Once identified, the two atom groups connected via the abnormal bond are translated along a vector away from the backbone of the residue. The system is subsequently minimised to correct any distortions and reassessed (Fig. 2B).

#### Equilibration

There are a number of levels of equilibration implemented by CG2AT2, depending on the required output. In the simplest and fastest instance, a non-equilibrated atomistic system may be produced by using the flag `-o none`. This means that the system is only energy minimised as part of the conversion, following the reintegration of the complete system. Beyond this, the default level of equilibration for CG2AT2 is to relax the system with a 5 ps NVT simulation with position restraints applied to the backbone Cα atoms of the protein, if included.

### Protein conversion types

To establish the atomistic protein coordinates, CG2AT2 enables both a *de novo* protocol, where the atoms are constructed from the CG beads, or an aligned approach, where a supplied atomistic protein structure is fitted with the CG protein coordinates. The decision of which approach to use depends upon whether one wishes to retain any protein conformational changes that have arisen from the CG simulation. The *de novo* method is also well suited to retaining protein interactions with non-protein groups such as lipids and minor conformational changes. Conversely, if particular atomic interactions are important, e.g. the orientation of the backbone within the selectivity filter of a potassium channel, or if the CG simulation is being used to establish an optimal bilayer as a starting point for an atomistic simulation, the aligned method should be used. CG2AT2 allows for a range of conversion types for the user, the choice of which to use depends upon the hypotheses being addressed by the simulation. Upon successful conversion, a root mean square deviation (RMSD) value comparing the backbone of each CG monomer to the converted atomistic protein is provided, as well as a summary of the atomistic system.

The *de novo* method is reliant on the coordinates of the CG beads. To improve the secondary structure, an additional correction applied to the amide bond, after the fragment alignment. In this correction the amide bond is aligned to the cross vector of the two proceeding backbone beads, as originally outlined by Wassenaar *et al*^13^. This enables the majority of the backbone hydrogen bond network to be recovered. Conversely, if a higher degree of similarity to the atomistic input structure is required, the backbone hydrogen bond network within the supplied atomistic structure is mapped. This network is used as distance restraints to guide the *de novo* NVT equilibration and thereby aid in the recovery of the secondary structure lost during the CG simulation. This allows the production of either an independent *de novo* or a guided *de novo* structure. The latter may be implemented by using the flag `-disre` and an input `-a` atomistic protein coordinate file (Fig. S1).

The aligned method requires a supplied atomistic structure, using the `-a` flag, and builds upon the *de novo* conversion. As with the *de novo* method the user is able to select the degree of information taken from the CG system. CG2AT2 contains four alignment methods for the supplied atomistic chains: individual atomistic chains (default), user-defined grouped atomistic chains, whole atomistic protein or grouped based on the CG subunits (Fig. 2C). Initially, the supplied structure is separated into individual chains, identified by a greater than 4 Å amide bond length. The amino acid sequence of each atomistic chain is then indexed to the CG protein sequence. Depending upon the choice of grouping, the atomistic backbone is fitted to the CG model. If only a partial structure is detected during the sequence alignment, CG2AT2 implements a hybrid build approach, i.e. combining *de novo* and aligned protein structures (Fig. 2D). In this instance, any residues missing from the input atomistic structure but within the CG protein, are incorporated from the *de novo* method. This is particularly useful for adding flexible, unfolded residues to a protein, e.g. growing residues onto a core CG protein to add lipoprotein tethers^23^.

The energy minimised *de novo* non-hydrogen atoms are steered onto the aligned protein coordinates generated by the aforementioned structural alignment. The steering is undertaken in a multi-step process of increasing harmonic restraints for 2 ps each (100 to 10,000 kJ mol^−1^ nm^−2^). This methodology preserves interactions between protein and non-protein residues with minimal perturbation, and compares favourably with other embedding methodologies, such g_membed, alchembed, inflategro and the original CG2AT^16,24–26^, in retaining protein-lipid interactions from CG. To minimise non-protein molecule displacement during the steered MD, an additional positional restraint is applied to the non-protein, non-hydrogen atoms.

### Macromolecular Systems

We tested CG2AT2 upon a range of increasingly complex systems from plain lipids bilayers to multi-membrane spanning protein complexes. The systems were all taken from ongoing projects with the research group using a mixture of Martini versions 2.0^27^, 2.2^28,29^ and 3.0 (beta). For all CG Martini simulations an elastic network of 1000 kJ mol^−1^ nm^−2^ was applied between all backbone beads between 0.5 and 1 nm. Electrostatics were described using the reaction field method, with a cut-off of 1.1 nm using the potential shift modifier and the van der Waals interactions were shifted between 0.9-1.1 nm. The temperature and pressure were kept constant throughout the simulation at 310 K and 1 bar respectively, with protein, lipids and water/ions coupled individually to a temperature bath by the V-rescale method^30^ and a semi-isotropic Parrinello-Rahman barostat^31^. Electrostatics were described using PME, with a cut-off of 1.2 nm and the van der Waals interactions were shifted between 1-1.2 nm. The TIP3P water model was used, the water bond angles and distances were constrained by SETTLE^32^. All other bonds were constrained using the LINCS algorithm^33^. The production simulations were run in an NPT ensemble with temperature V-rescale coupling at 310 K with protein, lipids and water/ions coupled individually and semi-isotropic Parrinello-Rahman barostat at 1 bar. All CGMD systems contain a salt concentration of 0.15 M NaCl. After 1 μs of CGMD simulation the CG systems were converted to a CHARMM36 atomistic form. To access the stability of the CG2AT conversions, the systems were equilibrated for 1 ns with 1000 kJ mol^−1^ nm^2^ position restraints applied to the Cα atoms after their conversion. A further 100 ns unrestrained MD simulation was then performed on each atomistic system. The following systems are ordered by system size from small (~37,000 atoms) to large (>2,000,000 atoms).

#### System 1 - A POPC membrane

A CG bilayer of 128 POPC lipids, solvated with 1,682 Martini water beads and a total system size of 3,256 beads. In comparison the converted atomistic system has 37,830 atoms.

#### System 2 - A mixed-lipid bilayer

A bilayer consisting of 80 POPC, 10 palmitoyl-oleoyl-phosphatidylserine (POPS) and 32 Cholesterol (CHOL) molecules was solvated with 1,778 Martini water beads, creating a total system size of 3,170 beads. The converted atomistic system has 36,362 atoms.

#### System 3 - SRY

The Sex-determining Region Y (SRY) protein recognises and binds to double stranded nucleotides. We used the *H. sapiens* structure (PDB ID: 1J46^34^) to create the CG representation and for CG2AT2 conversion. The system consists of a CG tetrameric SRY with 13 DNA base pairs, solvated with 18,661 Martini waters beads, yielding a system size of 28,938 beads. The atomistic system contains 324,468 atoms.

#### System 4 - δ-opioid

The class A G-protein coupled receptor (δ-opioid) activates the β-arrestin and G-protein pathways in the central nervous system. We used the *H. sapiens* structure (PDB ID: 4N6H^35^) to create the CG representation and for CG2AT2 conversion. The system consists of a CG δ-opioid, 237 POPC solvated with 3,743 Martini water beads yielding a system size of 7,338 beads. The converted atomistic system contains 82,569 atoms

#### System 5 - Lgt

The lipoprotein diacylglyceryl transferase (Lgt) transfers the diacylglyceryl group from a phospholipid to the thiol group of a conserved cysteine residue of the prolipoprotein. We used the *E. coli* model (PDB ID: 5AZC^36^) to create the CG representation and for CG2AT2 conversion. The system consists of a CG Lgt, 74 palmitoyl-oleoyl-phosphatidylglycerol (POPG) and 290 palmitoyl-oleoyl-phosphatidylethanolamine (POPE) solvated with 5,926 Martini waters beads yielding a system size of 11,674 beads. The converted atomistic system contains 126,028 atoms.

#### System 6 - LspA

The lipoprotein signal peptidase (LspA) catalyses the removal of signal peptides from prolipoproteins. We used the *S. aureus* model (PDB ID: 6RYO^37^) to create the CG representation and for CG2AT2 conversion. The system consists of a CG LspA, 74 POPG and 290 POPE solvated with 7,399 Martini waters beads, producing a system size of 12,928 beads. The system was converted to an atomistic system of 142,690 atoms.

#### System 7 - Lnt

The lipoprotein N-acyl transferase (Lnt) catalyses the phospholipid dependent N-acylation of the N-terminal cysteine of apolipoproteins. We used the *E. coli* model (PDB ID: 5N6H^38^) to create the CG representation and for CG2AT2 conversion. The system consists of a CG Lnt with 75 POPG and 288 POPE, solvated with 7,840 Martini waters beads and a total system size of 14,091 beads. The converted atomistic system has 153,299 atoms.

#### System 8 - LolB

This protein is an outer membrane lipoprotein receptor and receives a mature lipoprotein from LolA for delivery to the outer membrane of Gram-negative bacteria. We used the *E. coli* model (PDB ID: 1IWM^39^) to create the CG representation and for CG2AT2 conversion. The system consists of a CG LolB with 157 POPG and 363 POPE solvated with 9,736 Martini waters beads with a salt concentration of 0.15 M NaCl yielding a system size of 17,529 beads. The atomistic system consists of 193,138 atoms.

#### System 9 - Kir2.2

Kir2.2 is an inward rectifying potassium channel. We used the *G. gallus* model (PDB ID: 3SPI^40^) to create the CG representation and for CG2AT2 conversion. The system consists of a CG tetrameric Kir2.2 with 4 phosphatidylinositol-4,5-bisphosphate (PIP2) and 465 POPC solvated with 14,960 Martini waters beads and a total system size of 24,272 beads. The atomistic system contains 272,230 atoms.

#### System 10 - Wzm-Wzt

Wzm-Wzt transports fully elongated, lipid-linked lipopolysaccharide O-antigens across the inner membrane, using an undecaprenyl-linked carrier. We used the *A. aeolicus* structure (PDB ID: 6M96^41^) to create the CG representation and for CG2AT2 conversion. The system consists of a CG tetrameric Wzm-Wzt with 125 POPG and 504 POPE solvated with 18,661 Martini waters beads and therefore a system size of 28,938 beads. The atomistic system contains 324,468 atoms.

#### System 11 - Lpt complex

The lipopolysaccharide transport (Lpt) protein complex spans the periplasm of Gram-negative bacteria and shuttles lipopolysaccharide (LPS) molecules from the inner to outer membrane. We used *E. coli* structures and homology models (LptA: 2R19^42^, LptC: 3MY2^43^, LptB2FG: Swiss-model^44^ based on 6MIT^45^, LptDE: Swiss-model based on 4Q35 ^46^) to create the CG representation and for CG2AT2 conversion. The system consists of a CG Lpt complex composed of 4 LptA, 2 LptB, 1 LptC, 1 LptD, 1 LptE, 1 LptG and 1 LptF with 147 lipopolysaccharides, 284 POPG and 1,150 POPE solvated with 133,459 Martini waters beads with a total CG system size of 181,263 beads. The atomistic system has 2,039,940 atoms.

## Results

To assess the quality of the CG to atomistic conversion, CG2AT2 was evaluated against 11 macromolecular systems of varying size and complexity. To demonstrate the initial accuracy, RMSDs were calculated for each *de novo* endpoint against the input CG structure, this yielded excellent conversion RMSDs of less than 0.09 nm. The RMSD between the aligned structures to the CG demonstrates the fluctuations of the CG structure over time. Therefore, the RMSD between the aligned and the supplied atomistic structures was measured. This yielded RMSDs of less than 0.013 nm, within range of a standard minimised system. These results are summarised within Table 1.

**Table 1.** The similarities between the Martini backbone and Cα atom of each residue after End 1 and 2. The final RMSDs of the Cα atoms after 100 ns atomistic simulation are reported.

| System | Protein monomers | *de novo* vs CG |  | *de novo* NVT + 100 ns sim | Aligned vs CG | Aligned vs input AT | Aligned + 100 ns sim |
| --- | --- | --- | --- | --- | --- | --- | --- |
|  |  | MIN | NVT |  |  |  |  |
| **SRY (3)** | 1 | 0.072 | 0.093 | 0.358 | 0.352 | 0.012 | 0.196 |
| **δ-opioid (4)** | 1 | 0.089 | 0.102 | 0.266 | 0.162 | 0.009 | 0.212 |
| **Lgt (5)** | 1 | 0.090 | 0.164 | 0.323 | 0.192 | 0.010 | 0.255 |
| **LspA (6)** | 1 | 0.088 | 0.150 | 0.293 | 0.159 | 0.011 | 0.230 |
| **Lnt (7)** | 1 | 0.088 | 0.127 | 0.183 | 0.098 | 0.010 | 0.198 |
| **LolB (8)** | 1 | 0.088 | 0.154 | 0.209 | 0.181 | 0.011 | 0.369 |
| **Kir2.2 (9)** | 4 | 0.087 | 0.105 | 0.386 | 0.152 | 0.010 | 0.261 |
| **WzmWzt (10)** | 4 | 0.087 | 0.147 | 0.329 | 0.187 | 0.013 | 0.225 |
| **Lpt (11)** | 11 | 0.090 | 0.217 | n/a | 0.305 | 0.011 | n/a |

For CG2AT2 to be included in a workflow, the conversion must ideally be undertaken in a high-throughput manner. Whilst, the conversion time is highly dependent upon the specifications of the machine used to run CG2AT2, we provide a set of benchmarks as a guide for the user. Figure 3 shows the cumulative time required for each step in the conversion with the possible endpoints for the conversion are shown. CG2AT2 takes roughly 4 minutes for a full conversion of a 100 K atom system producing an equilibrated *de novo* and aligned system. As expected, the majority of the time taken to convert the system is the Molecular Dynamics equilibration steps namely the NVT and the multi-step steering.

**Figure 3:**
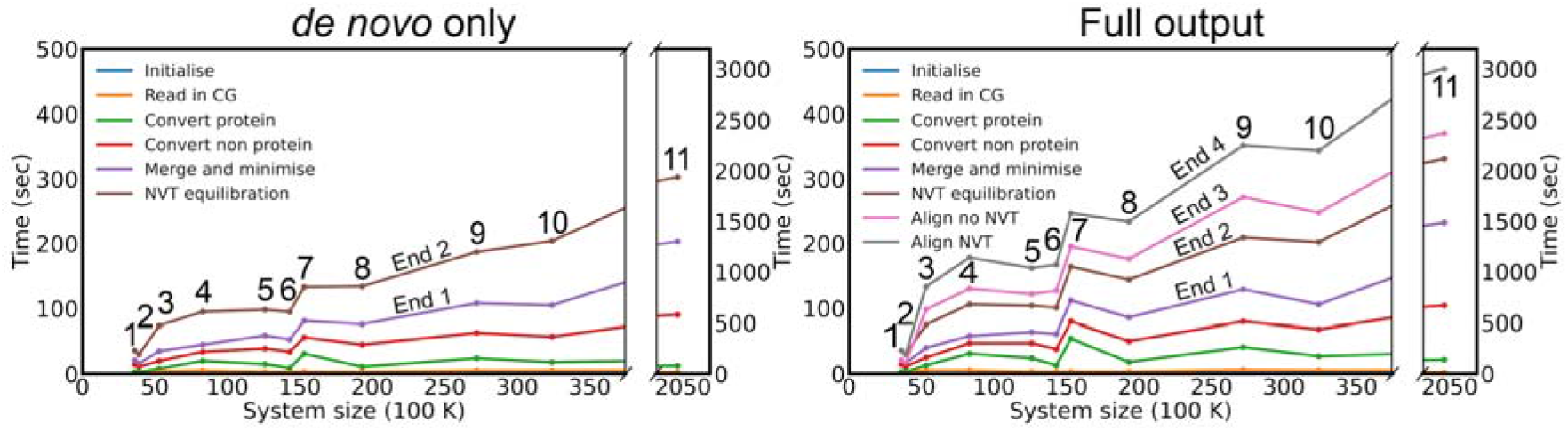
The timings for the conversion from CG to the CHARMM36 atomistic representation. The data shows the cumulative time plotted against the final atomistic size of each system. End 1 corresponds to the minimised *de novo* system, End 2 corresponds to the *de novo* after which a short NVT simulation has been undertaken, End 3 corresponds to the production of the aligned structure without NVT equilibration and End 4 corresponds to the production of the aligned structure with an NVT equilibration. The conversion was run on a i7-6800K intel processor, GeForce GTX 1080 GPU and 64 Gb of RAM using GROMACS 2020 with the parallelisation of CG2AT2 limited to 8 threads.

### Solvent

A significant update to CG2AT now allows for the full conversion of the CG systems, including waters and ions. Within the new implementation of solvent conversion, CG2AT2 places a randomly rotated arrangement of solvent molecules over each CG solvent bead. The solvent fragment can contain any number of molecules in any configuration. Here, we utilise a fragment containing four water molecules in a roughly tetrahedral arrangement taken from an equilibrated atomistic MD system. Additional solvent molecules/models can easily be added as additional solvent fragments. As CG ions are treated as solvated, the solvent fragment is overlaid over the CG ion coordinate. This process is demonstrated within Figure 4. The similarity of the CG to AT solvent bulk properties is demonstrated in Figure 4C. Here, we see similar density profiles in the complex membrane before and after conversion. To assess the quality of the solvent conversion, we analysed the radial distribution function (RDF) of the water oxygen atom against Na^+^, Cl^−^ and itself (Fig. 4D). The quality of the initial solvent conversion is demonstrated by the minimal shift in the peak RDF values of less than 0.014 nm between initial minimisation and the equilibrium values, demonstrating the need for minimal equilibration post conversion.

**Figure 4:**
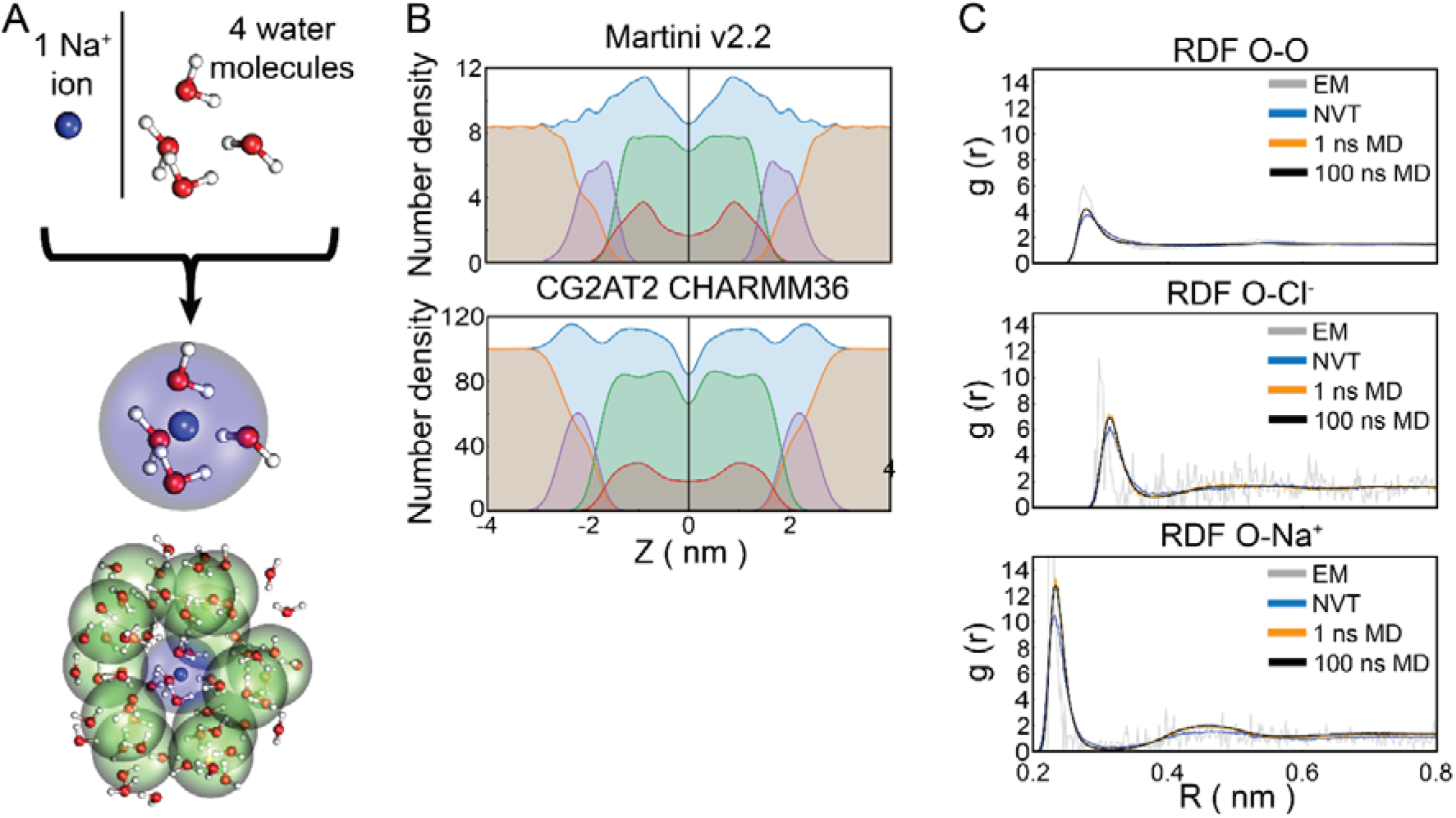
Solvent conversion. (**A**) Ion conversion; A tetrahedral arrangement of water is placed over a single ion located at the centre of the CG ion bead. A zoomed-out view of a Na^+^ ion surrounded by a cluster of CG waters. (**C**) Number density profiles of the complex membrane system. Colours indicate: System (blue), water and ions (orange), sterols (red), phospholipid tails (green) and the phospholipid headgroup (purple). (**D**) O, Cl^−^ and Na^+^ atom RDFs referenced to the TIP3P oxygen atom, EM: Complete system energy minimisation by the steepest descent algorithm. NVT: Optional unrestrained 5 ps NVT equilibration. 1 ns MD: An initial 1 ns position restrained NPT equilibration. 100 ns MD: A 100 ns NPT production run.

### Lipids

An advantage of converting CG simulations to atomistic representation is the retention of the equilibrated bilayer properties, such correctly mixed fluid-phase lipid bilayer. We tested a conversion of a complex bilayer composed of POPC, POPS and CHOL at a ratio of 8:1:3 solvated with 0.15 M NaCl. An identical system was created by the webserver CHARMM-GUI as a direct comparison^47^. Both systems were equilibrated for 1 ns NPT before production runs. The area per lipid (APL) was compared between the CG, CG2AT2 and CHARMM-GUI. Between the CG2AT2 conversion and CHARMM-GUI membrane, the APL of the lipid components are in excellent agreement (Fig S2A). We also calculated the lipid tail order parameters profiles for the AT systems for both bilayers produced by CG2AT2 and CHARMM-GUI (Fig. S2B). After 100 ns the order parameters of both sets of simulations closely match. Interestingly however, the rate of convergence to the final profile varies between the two methods, with CG2AT2 converging to a mean Δ-S_CH_ at less than 0.005 roughly ~25 ns faster than CHARMM-GUI (Fig. S2C).

### Assessing the quality of the conversion by evaluating protein dynamics

We employed a set of nine protein test systems, spanning a range of secondary structure complexities and architectures. These systems were used to evaluate the conversion procedure of CG2AT2 in terms of the stability and quality of the dynamics over a subsequent 100 ns of atomistic simulation post-conversion. In all cases, the majority of the secondary structure elements were recovered from the CG simulations (Fig. 5).

LolB was used to demonstrate the hybrid conversion approach (Fig. 2D). The β-barrel core of the LolB protein was solved by X-ray crystallography, however this lacks the N-terminal region, which is anchored to the membrane by a triacyl-cysteine. The N-terminal domain and the lipoprotein tether were then grown onto the β-barrel in CG representation using the method outlined in Rao, S. *et al*^23^.

To measure the similarity of the CG and the converted systems, structures were compared using Cα and BB RMSDs. A summary of conversion RMSDs is shown in Table 1. Depending on the level of equilibration chosen, there is an increasing degree of conformational drift. The *de novo* conversion supplies a system closest to the initial CG representation, with an post-minimisation RMSD of 0.087 ± 0.005 nm compared to 0.140 ± 0.036 nm after NVT equilibration. Whilst a comparison between the aligned protein structure and the CG model can be made, this mostly highlights the deviation of the CG conformation from the input atomistic structure. A more useful measurement is between the aligned structure generated by the steered MD step and the input atomistic structure. This typically provides an RMSD of 0.011 ± 0.002 nm highlighting that by using the steered MD, the supplied structure can be accurately reproduced. By applying variations in the aligned RMSD, any clashes between aligned monomers and/or deviations arising due to the hybridisation method can be easily detected and highlighted by CG2AT2 (0.015 nm cut-off).

The backbone provides much of the structural stability of the protein. Therefore, we calculated a Ramachandran plot to assess the stereochemical quality of the *de novo* and aligned output of CG2AT2 (Fig. S3). As a guide, a good atomic resolution protein model will generally expect over 90 % of residues within the allowed regions^48^. Here, the *de novo* conversion shows an excellent level of quality with 93-99% within the allowed regions with the majority of the test systems above 96 %. This is also in excellent agreement to the aligned structures, which were resolved at the atomic resolution.

Whilst the RMSD and Ramachandran plots demonstrate the accuracy and quality of the conversion, they provide little information on the stability of the systems. Therefore, the converted systems were simulated for 100 ns atomistic MD simulations using the standard CHARMM36 force field settings. The subsequent conformational drift of the protein was analysed by RMSD, root mean square fluctuation (RMSF) and secondary structure retention (Table 1, Fig. 5 and Table S1, Fig. S4). In all cases, the structures remained stable throughout the 100 ns simulation. The global deviation of the simulations from their initial configuration is highlighted by the RMSD. The *de novo* simulations demonstrated a slightly larger degree of conformational drift over the course of the simulation with an average RMSD of 0.293 ± 0.066 nm compared to 0.243 ± 0.052 nm for the aligned systems. To pinpoint areas of instability and dynamic regions of the converted system, we evaluated the RMSF calculations. The nine test systems shown here demonstrate comparable RMSFs to the aligned systems, with only the extended loops and short helices showing values greater than 0.2 nm. This is most noticeable within the periplasmic region of Lgt and the N- and C-terminal regions of Kir2.2.

Whilst the tertiary and secondary structures within the Martini simulations are retained via the use of elastic networks, minor changes can still occur within the protein structure. We therefore expect to see the majority of secondary structure to be recovered during the *de novo* conversion, as detailed in the original atomistic protein coordinate file. A qualitative comparison of the secondary structures in Figure 5, demonstrates that a large degree of secondary structure is recovered and retained over the 100 ns simulation. There are also only minor variations in the secondary structure retention, when using the *de novo* approach, and this is in excellent agreement with the aligned systems. This is further highlighted by the Jaccard indices, which represent the secondary structure correlation to a reference structure. The numbers presented here represent the mean of the simulations means. The aligned structures were used as a guide to estimate the secondary structure variability over 100 ns of atomistic simulation, with a mean index of 0.81 ± 0.07. A Jaccard index was calculated for the *de novo* conversion in relation to the input structure to test the ability of CG2AT2 to recover the original secondary structure, with a mean index of 0.69 ± 0.09. A Jaccard index of 0.72 ± 0.07 was also calculated to compare the *de novo* simulation to the *de novo* starting structure. These values demonstrate that the dynamics of the *de novo* conversion are marginally higher than the aligned systems. This appears to arise from drift in the β-strands within the CG system. Recovery of short helices as can be seen within Figure 5, in particular the LspA and Lgt *de novo* conversions.

**Figure 5:**
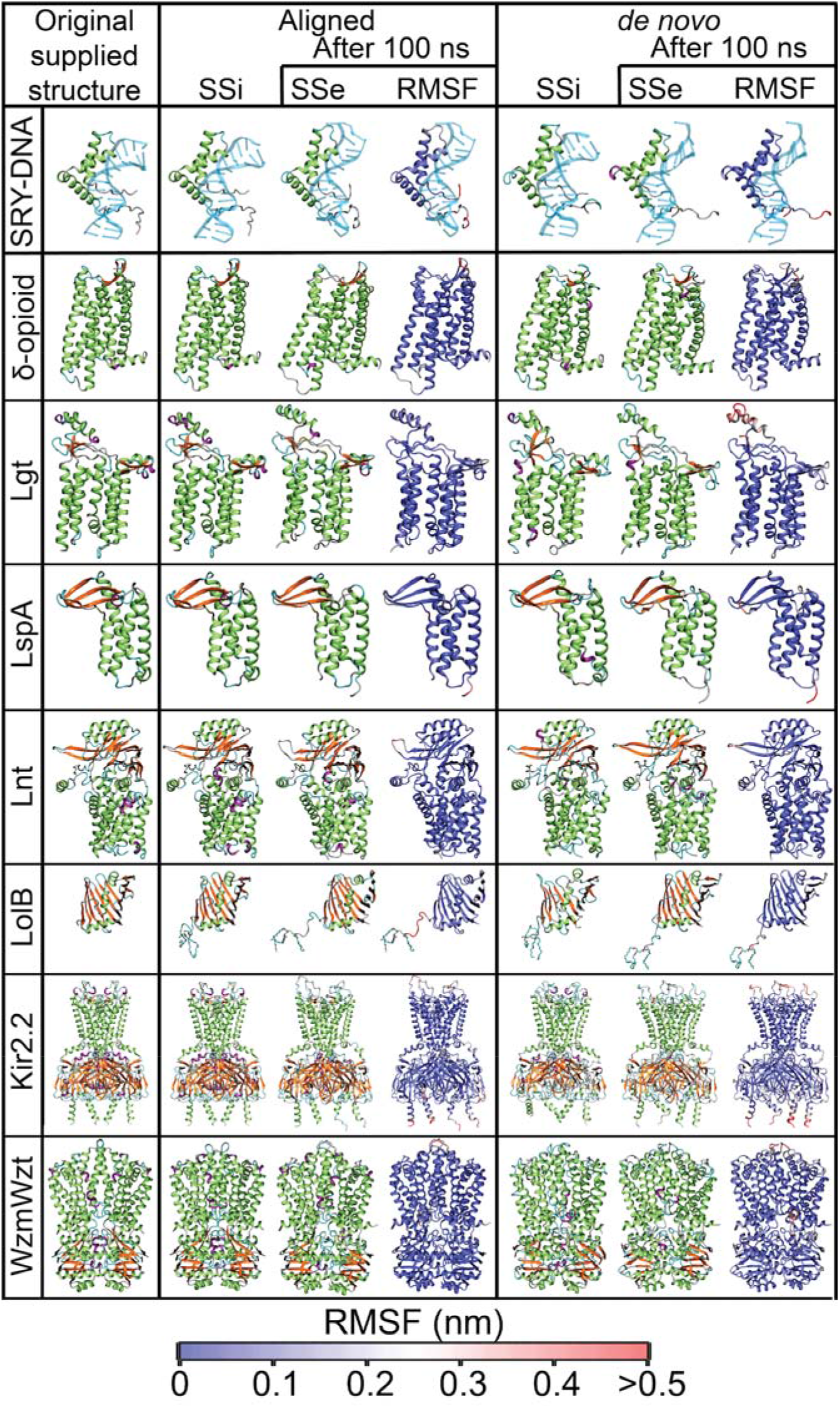
Protein secondary structure and stability. Original supplied structures from the RCSB database. (SSi) The protein structure after initial conversion colour by secondary structure. (SSe) The protein secondary structure after 100 ns of atomistic simulation, coloured by the end secondary structure. (RMSF) The protein structure after 100 ns coloured by the RMSF, on a blue to red scale, with each monomer aligned individually. The colour scheme used for the secondary structure is as follows. Turn: cyan, Extended sheet: orange, Bridge: tan, α-helix: lime, 3_10_-helix: purple, π-helix: blue and coil: silver.

### Protein-ligand interactions

Two of the core advantages of CG simulations are straightforward system configuration, e.g. *insane*, and quick equilibration. A number of papers have demonstrated the use of CG simulations to find difficult small molecule ligand coordination sites, lipid binding sites and protein-protein interactions^49^. Here, we demonstrate the conversion of the tetrameric Kir2.2 protein coordinated with PIP_2_ lipids. To generate a Kir2.2:PIP_2_ bound pose, the system was simulated for 1 μs. The final frame from the CG simulation captured two PIP2 molecules bound, as described previously (Corey *et al*^6^). We show qualitatively that the steering process to generate the aligned systems minimally perturbs the binding pattern between the lipid and the protein (Fig. 6). The RMSD between the *de novo* and aligned PIP_2_ non-hydrogen atoms shown in binding sites 1 and 2, are 0.25 and 0.29 nm respectively.

**Figure 6:**
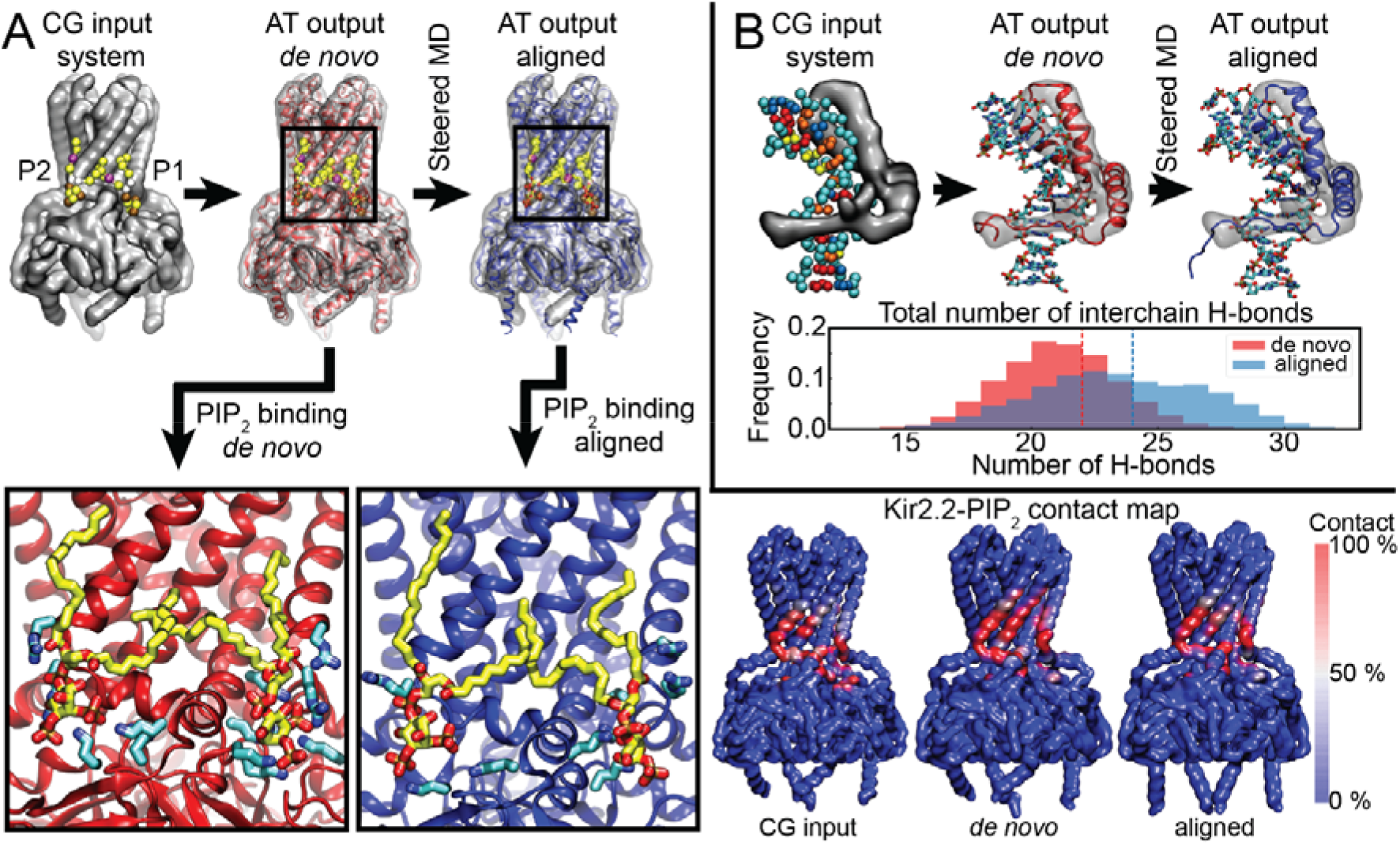
Protein: Ligand conversion and retention. (**A**) Conversion of the Kir2.2 bound to four PIP_2_ lipids. Shown here are two PIP_2_ lipids bound in pose 1 and 2. The PIP_2_ binding sites found in the CG simulations are retained upon both *de novo* and aligned conversions. (bottom right) The contact maps of PIP2 binding to Kir2.2. The last 100 ns of the CG simulation is shown as a comparison with the 100 ns atomistic simulations. (**B**) Conversion of the SRY protein in complex with double stranded DNA. The number of DNA interchain hydrogen bonds over the 100 ns are shown, the dotted line highlights the starting number of hydrogen bonds.

The increased RMSD arises from the inevitable rearrangement of amino acids leading to small steric clashes and rearrangement of electrostatic interactions between the aligned protein and PIP2. To test the stability of the PIP_2_ binding pose, the lipid contact maps of the converted systems were compared to the CG system. To provide a comparable map, the contacts within the last 100 ns of the CG simulation were compared to the 100 ns atomistic simulation. As shown by Figure 6, the contact maps between the three simulations are near identical, demonstrating the preservation of the protein-ligand interface. The Pearson correlation coefficient between the converted atomistic simulations and the CG simulation provide values of 0.71 and 0.70 for the *de novo* and aligned systems, respectively. The high correlation of 0.77 between the *de novo* and aligned systems, demonstrates that little modification has occurred to the protein-PIP_2_ interactions as a result of the steered MD alignment.

Another complex system conversion problem are protein complexes that contain DNA. Here, we demonstrate the variability of CG2AT2 with the conversion of the SRY protein in a complex with 14 base pairs of double stranded DNA (Fig. 6B). The multimeric non protein residues are converted in a similar manner to the protein *de novo* conversion with the exception of the further processing of the protein backbone. To analyse the stability of the protein-DNA complex over the course of the simulations, the number of DNA interchain hydrogen bonds were calculated. We show in Figure 6B that the hydrogen bonding pattern remains stable in both *de novo* and aligned atomistic simulations with the total number of H-bonds being 29.0 ± 3.0 and 26.3 ± 2.5 respectively.

## Discussion

We have described an updated fragment-based approach of CG2AT for the accurate conversion of complex CG representations to atomistic detail. Here, we have demonstrated against a variety of systems that CG2AT2 can provide a gradient of conversion approaches from completely *de novo* to aligned, depending upon the degree of protein conformational changes in the CG system the user wishes to retain. A graphical representation of the workflow of CG2AT is demonstrated by the multicomponent CG Lpt system, composed of 11 protein monomers and two lipid bilayers (1,434 lipids and 147 LPS) (Figure 7).

**Figure 7:**
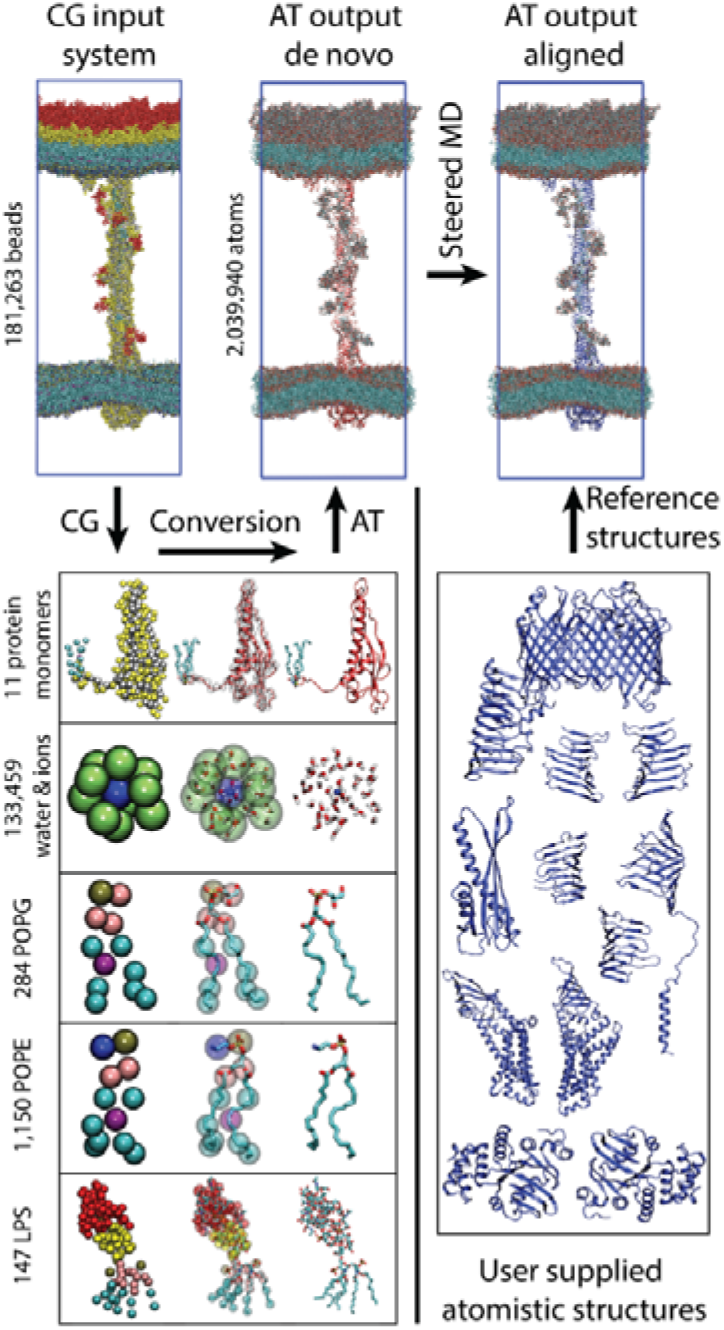
CG2AT2 conversion of the Lpt system to CHARMM36. The Martini CG system (Top left) is separated into individual residue types and then converted into their atomistic representation (Bottom left). The individual residue types are recombined providing the *de novo* system before being steered to the reference structures provided (Bottom right). For clarity, the solvent has been omitted from the images of the system.

CG2AT2 now performs complete reconstruction of atomistic systems from CG, including water. This is achieved almost entirely within python-based code, with GROMACS called to generate topologies and to run the equilibration simulations. CG2AT2 can convert complete systems with no pre-processing of the input files, via a supplied comprehensive database of over 107 unique residue types, at the time of publication. The current individual fragment libraries include standard CHARMM36, CHARMM36 with virtual sites and Stockholm lipids (Slipids)^50^ for conversion from CG Martini versions 2 and 3 (beta). To enable the variability of the residue types, we have included a range of updated force fields (AMBER99SB-ILDN with Slipids and CHARMM36) containing additional parameters as well as the base level force fields. However, it should be noted that all GROMACS readable force fields are compatible with CG2AT2.

The protocol outlined here generates the key files required for a production run simulation including an equilibrated atomistic system and topology files. This enables the user to implement high-throughput robust and accurate conversion of CG systems with minimal further processing. In combination with tools such as TS2CG^51^, CG2AT2 may enable conversion of triangulated surfaces all the way down to full atomic-level detail, and thereby permit flexible approaches for converting between molecular systems at multiple levels of granularity.

## Availability

CG2AT2 is available from github: https://github.com/owenvickery/cg2at, and may be added to a Conda installation using: conda install -c stansfeld_rg cg2at. For full usage instructions, please see https://github.com/owenvickery/cg2at.

## Acknowledgements

This work was supported by the Wellcome [208361/Z/17/Z], the BBSRC [BB/P01948X/1, BB/R002517/1 and BB/S003339/1] and MRC [MR/S009213/1]. ARCHER UK National Supercomputing Service (http://www.archer.ac.uk), provided by HECBioSim, the UK High End Computing Consortium for Biomolecular Simulation (hecbiosim.ac.uk), which is supported by the EPSRC (EP/L000253/1). We acknowledge the use of Athena at HPC Midlands+, which was funded by the EPSRC on grant EP/P020232/1, and the University of Warwick Scientific Computing Research Technology Platform for computational access. We thank Miriam Shiradski, Joshua Sauer, Michael Horrell, Robin Corey and Mark Sansom for useful discussions and testing of earlier versions of the methodology.

## Supplementary Information

**Figure S1:**
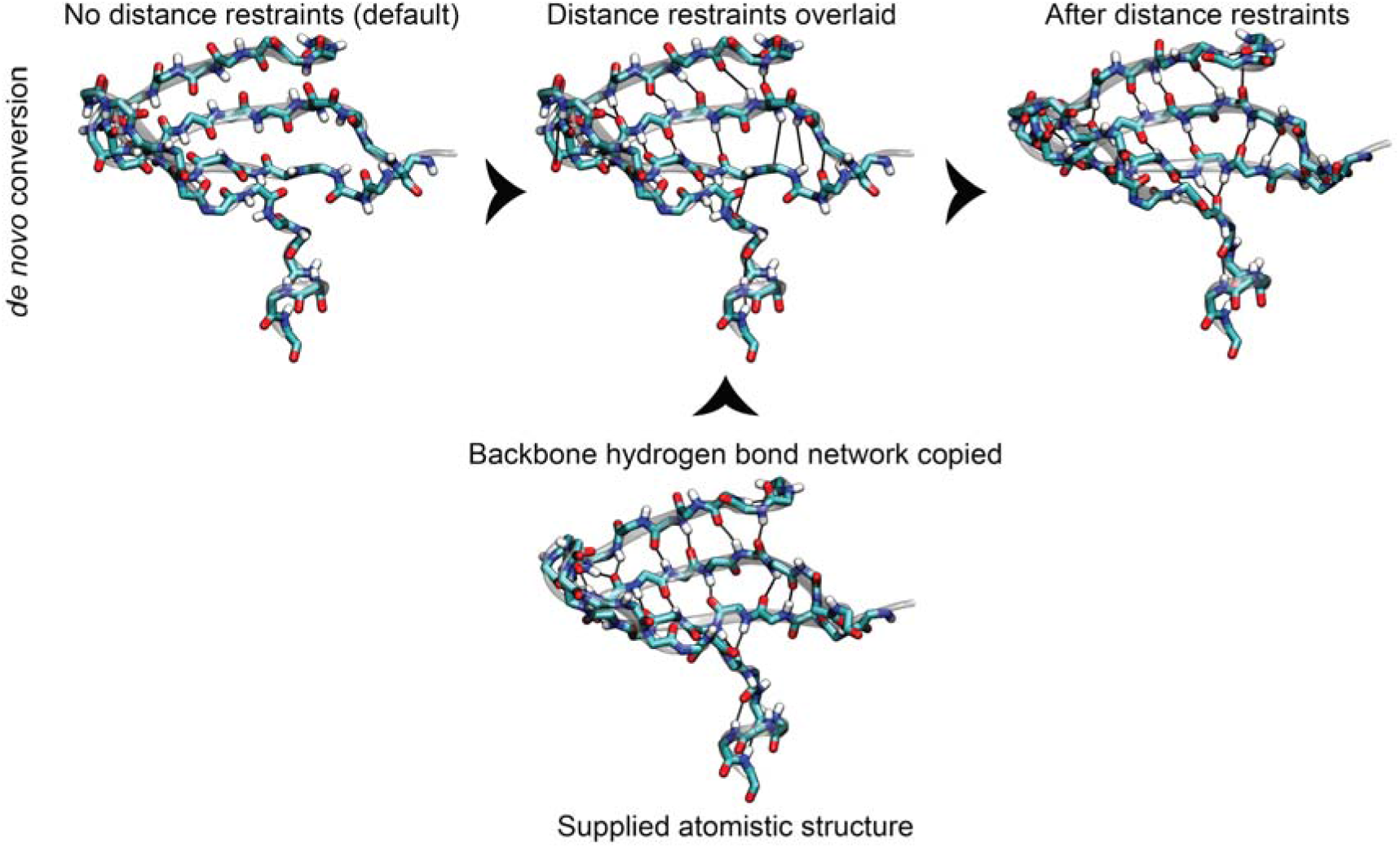
Backbone distance restraint matrix. Following the initial *de novo* conversion (Top left) an optional backbone distance restraint matrix can be applied. The backbone hydrogen bonds are mapped from the supplied atomistic structures (Bottom). The matrix is then applied to the *de novo* structures during the NVT equilibration (Top middle). This gives rise to a guided *de novo* conversion, containing the key *de novo* features with a corrected backbone.

**Figure S2:**
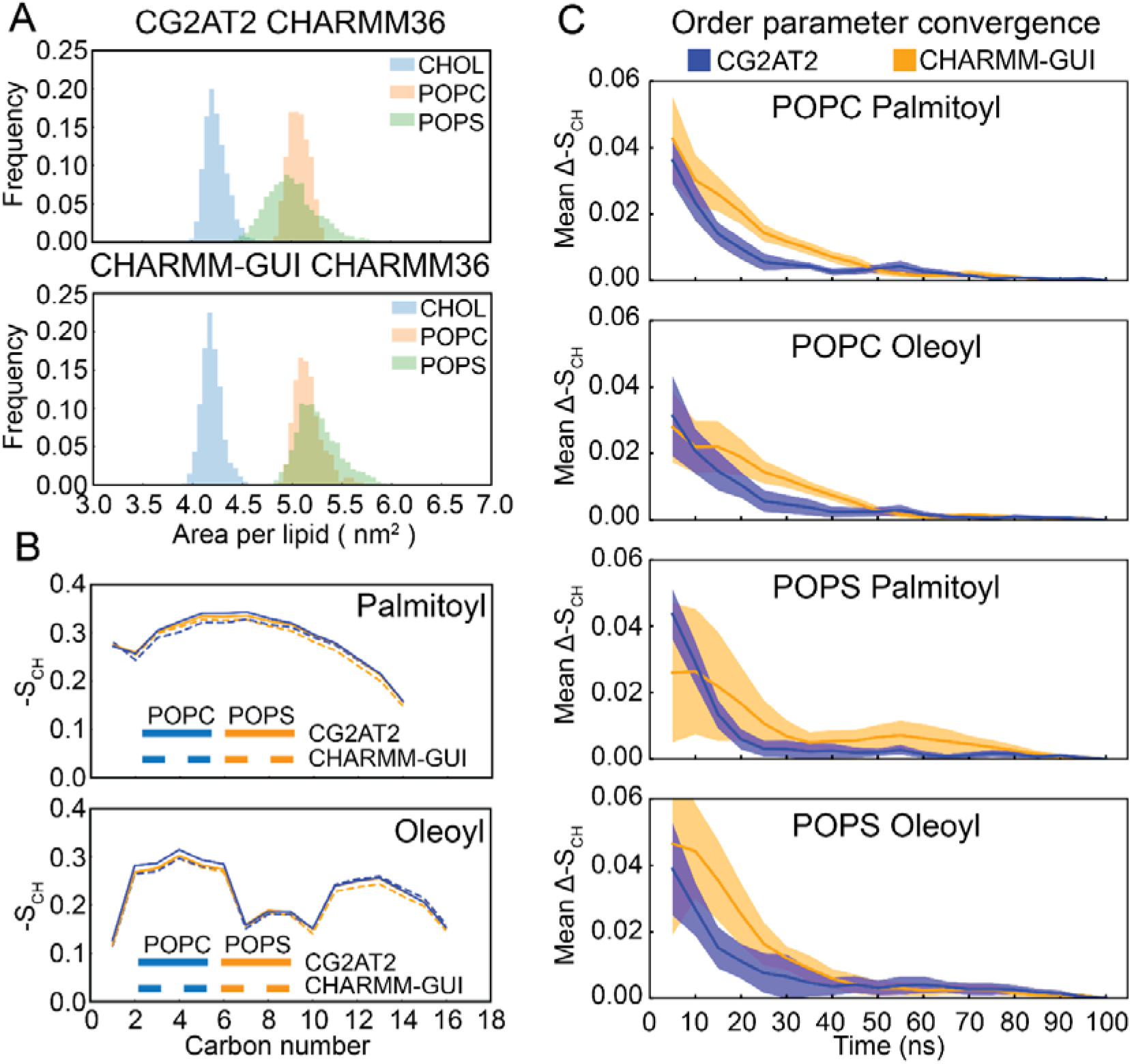
Lipid conversion. (**A**) A comparison of the APL of Martini v2.2, CG2AT2 and CHARMM-GUI. Dotted lines denote initial APL. (**B**) A comparison of deuterium order parameters of lipids between a system generated by CG2AT2 and the CHARMM-GUI webserver: POPC (blue) and POPS (orange). (**C**) The deuterium order parameters were calculated with increments of 5 ns up to 100 ns. The mean order parameter difference to the complete 100 ns is shown, shaded areas represent the standard deviation of order parameter difference.

**Figure S3:**
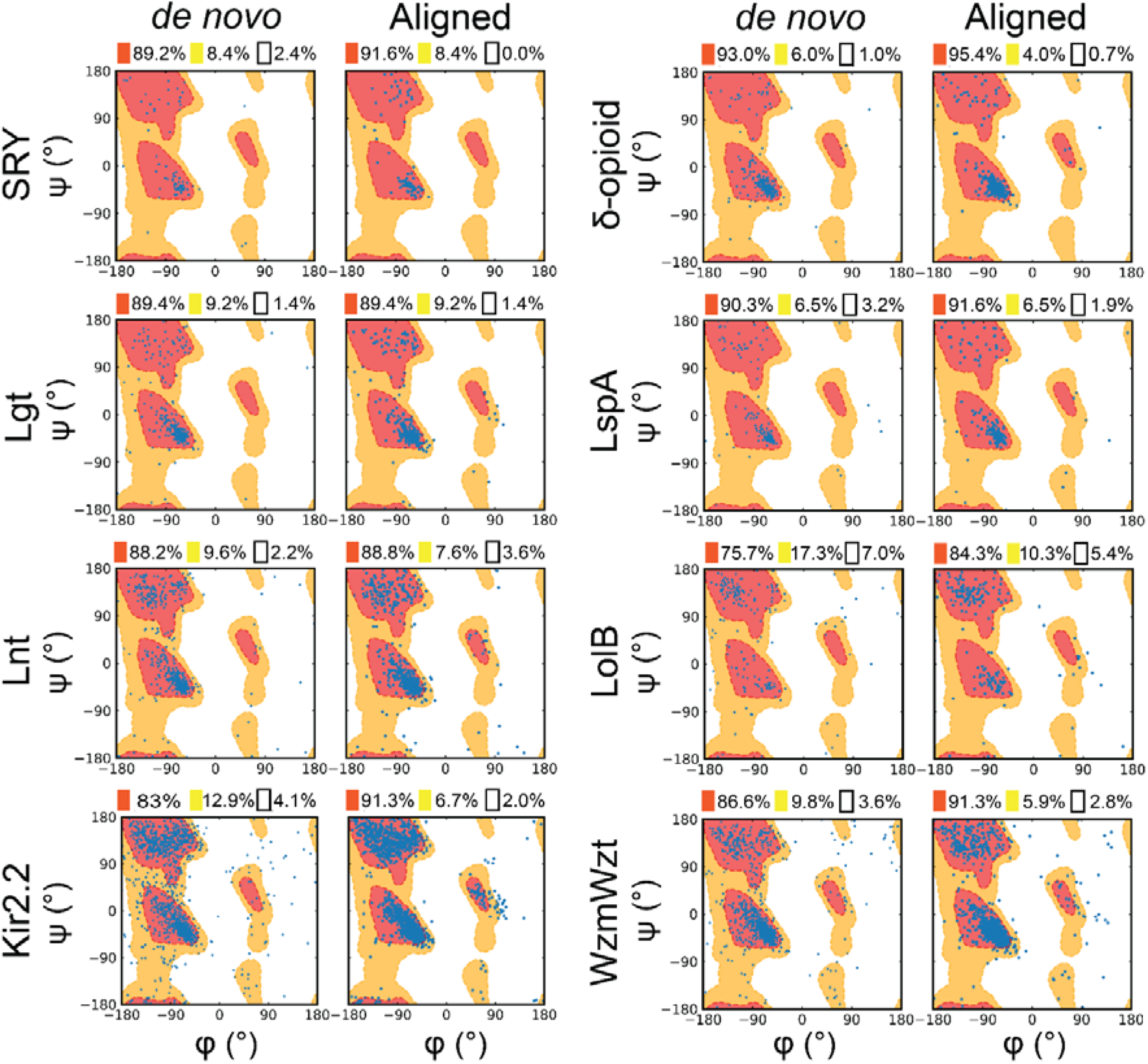
Ramachandran plots of converted systems. The *de novo* and aligned outputs of CG2AT2 were analysed using “gmx rama”. The conversions show an high level of quality with 93-99% in the allowed regions.

**Table S1:**
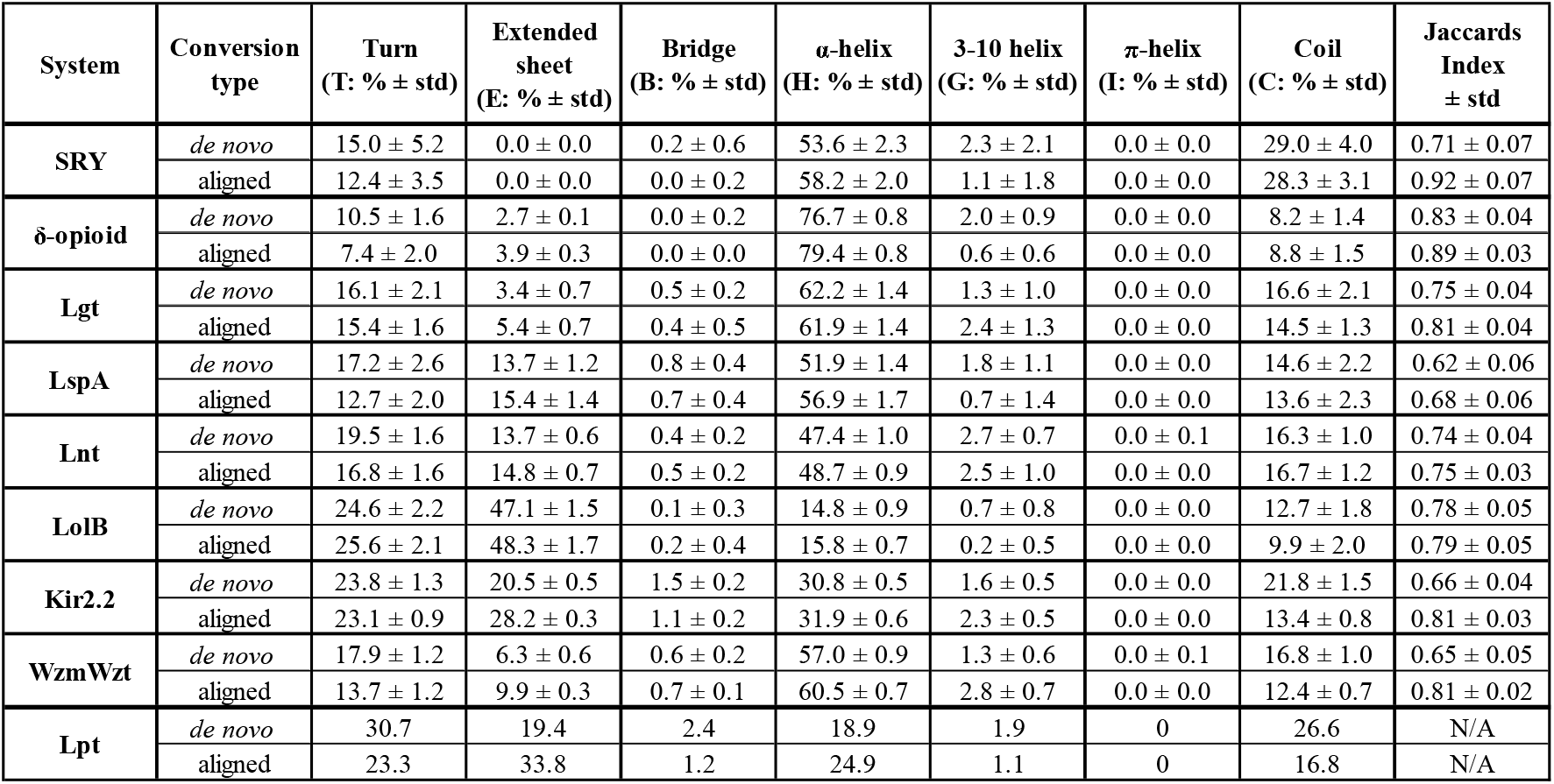
Secondary structure analysis of converted systems. The secondary structure was calculated every 1 ns over the 100 ns simulation. Shown here is the secondary structure composition breakdown for each conversion. The secondary structure similarity between the starting structure and each snapshot was calculated as a Jaccard index, J(A,B)= |A∩B|/|A⋃B|, the numerator is the number of residues which have the same secondary structure and the denominator is the total number of residues.

**Figure S4:**
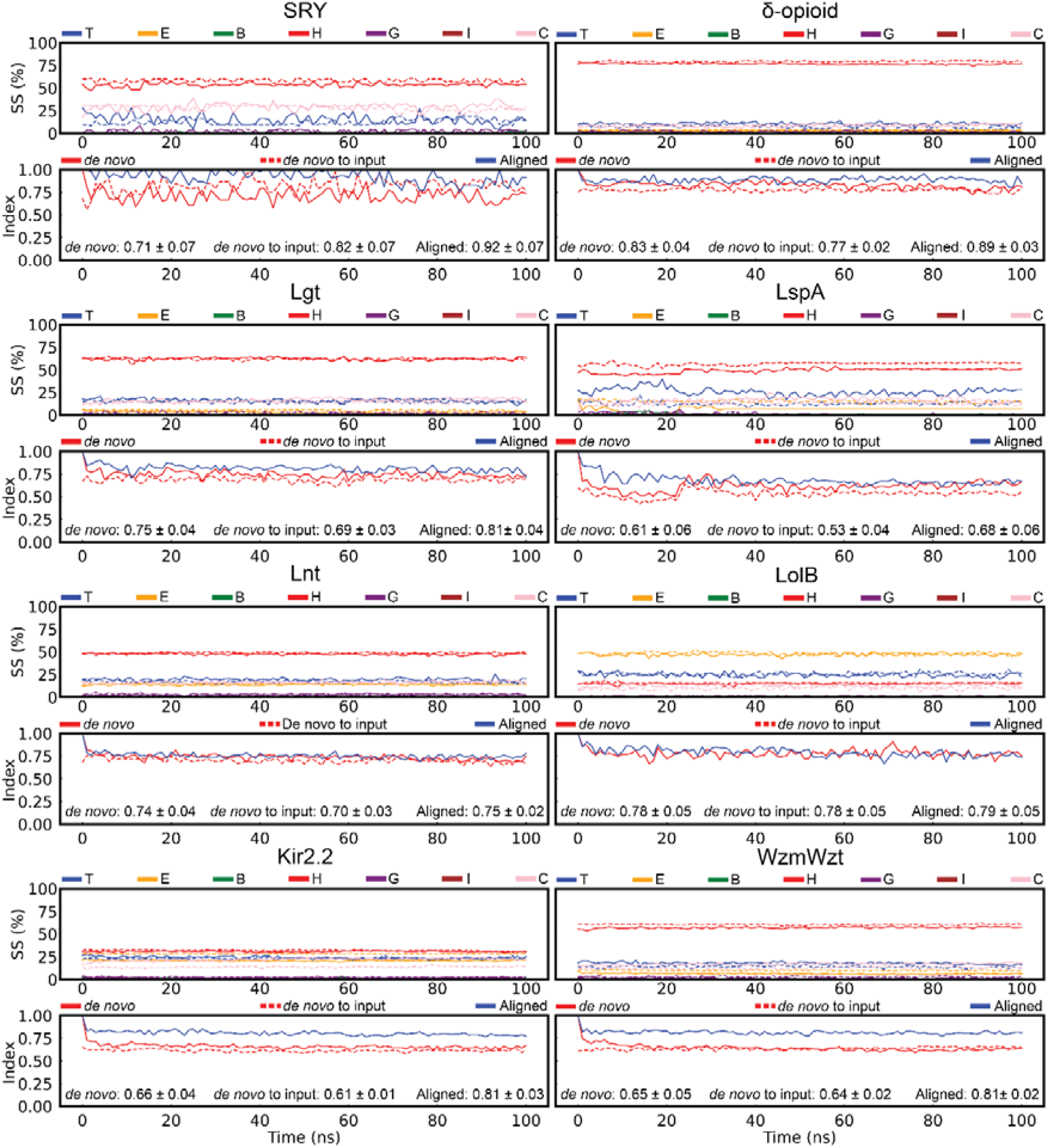
Secondary structure analysis. The secondary structure composition and Jaccard index shown for each 100 ns atomistic simulation. The scheme used for the secondary structure is as follows. T: Turn, E: Extended sheet, B: Bridge, H: α-helix, G: 3-10 helix, I: π-helix and C: Coil. Dotted lines for aligned and solid for *de novo*.

